# Nature lessons: the whitefly bacterial endosymbiont is a minimal amino acid factory with unusual energetics

**DOI:** 10.1101/043349

**Authors:** Jorge Calle-Espinosa, Miguel Ponce-de-Leon, Diego Santos-Garcia, Francisco J. Silva, Francisco Montero, Juli Peretó

## Abstract

Bacterial lineages that establish obligate symbiotic associations with insect hosts are known to possess highly reduced genomes with streamlined metabolic functions that are commonly focused on amino acid and vitamin synthesis. We constructed a genome-scale metabolic model of the whitefly bacterial endosymbiont *Candidatus* Portiera aleyrodidarum to study the energy production capabilities using stoichiometric analysis. Strikingly, the results suggest that the energetic metabolism of the bacterial endosymbiont relies on the use of pathways related to the synthesis of amino acids and carotenoids. A deeper insight showed that the ATP production via carotenoid synthesis may also have a potential role in the regulation of amino acid production. The coupling of energy production to anabolism suggest that minimization of metabolic networks as a consequence of genome size reduction does not necessarily limit the biosynthetic potential of obligate endosymbionts.

---

Nutritionally driven symbioses between insects and one or more intracellular, maternally inherited bacteria (endosymbionts) are widespread(1). These mutualistic associations allow insects to colonize novel ecological niches with unbalanced nutritional sources. The extreme difficulty to culture obligate endosymbionts outside of their hosts has limited the study of the interactions between symbionts and hosts mainly by using comparative genomics(2). Obligate endosymbionts usually have highly reduced genomes (<1,000 kb), which contain a conserved core of housekeeping genes, and genes for the biosynthesis of amino acids and vitamins that are essential for the host(1). The synthesis of these essential compounds requires, in many cases, the metabolic complementation with the host or from other coexisting endosymbionts(3). Although the nutritional role of obligate endosymbionts is well defined, other important aspects of the endosymbiont's biochemistry are less known. In spite of the fact that the metabolic networks of endosymbionts are relatively small, they are significantly complex and systems biology approaches are needed to further understand the physiology of these microorganisms. Earlier research approaches used techniques derived from graph theory and stoichiometric analysis. In particular, these techniques have been used to infer plausible scenarios of metabolic complementation(4-8), to determine the fragilities of the metabolic networks(5–8), to reveal characteristics of the transit from free-life to endosymbiosis(5, 8) and to further understanding of nitrogen management in these systems(5, 7).

However, to the best of our knowledge, it is still unclear to which extent endosymbionts are dependent upon their hosts for the necessary energy to support cellular processes and no previous work (neither theoretical nor experimental) has focused on this aspect of the endosymbiont's biology. Specifically, can these bacteria be energetically independent from their host? And, if this is the case, how do they operate to perform such a task? The answer to these questions is important not only for an understanding of insect-bacteria symbiosis, but also in order to set the limits of genome reduction properly, a matter of major importance in fields such as synthetic biology.

Among the known endosymbionts, *Candidatus* Portiera aleyrodidarum (hereafter referred to as *Portiera*), hosted in the whitefly bacteriocyte(9), displays a key characteristic that make it a good candidate for a study in this direction: it contains the set of genes coding for cytochrome oxidase, NADH dehydrogenase, and ATP synthase, and has the most reduced genome among endosymbionts whith these traits(10). This feature suggests that potentially *Portiera* is energetically autonomous even though the endosymbiont metabolic network, in obligate cooperation with the host, is involved in the synthesis of essential amino acids and carotenoids(11–15). The results presented in this paper support the hypothesis that *Portiera* is capable of synthesized its own ATP from the resources provided by the whitefly, due to the utilization of a limited number of pathways related to the synthesis of well-defined amino acids and/or carotenoids. Moreover, the involvement of the carotenoid synthesis in ATP production suggests that this pathway is a potential point for the insect to regulate amino acid production. Thus, the characterization of *Portiera’s* metabolism demonstrates the richness of the energetics of endosymbionts.

## Background

The structure of any metabolic network can be represented by its corresponding stoichiometric matrix ***N***_***mxn***_, where each element *n*_*ij*_ corresponds to the stoichiometric coefficient of metabolite *i* in reaction *j*. Hereafter, along with ***N***_***mxn***_ several constraints are needed to properly assess the properties of the associated metabolic network: the thermodynamic constraints, which prevents the irreversible reactions from taking negative flux values; the lower and upper bounds imposed over each reaction flux to represent specific environments (*e.g*. amount of available nutrients) or enzymatic/transporter maximum capacities; and the specific kinetic laws that govern each reaction of the network.

Unfortunately, the above-mentioned kinetic laws are often unknown. However, metabolism is characterized by usually fast reactions and high turnover of metabolites when compared to regulatory events and thus, a pseudo-stationary state (*i.e*. the metabolite concentrations are assumed constant) can be assumed as an approximation. Taking all restrictions into account, the set of all possible flux distributions (*i.e*. vector of reaction fluxes), the so-called flux space (F), is defined as follows:

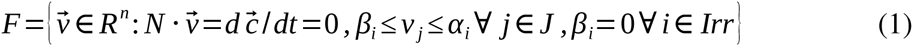

where *v* is the flux distribution, *c* is the vector of metabolite concentrations, *β*_*i*_ and *α*_*i*_ correspond to the lower and upper bound of reaction ***i***, respectively; and *Irr* is the set of the irreversible reaction indexes.

Elementary flux modes (EFM) are a special set of flux distributions which fulfills the constrains in equation (1) and is non-decomposable, that is, no reaction can be deleted from an EFM and still obtain a valid (non-trivial) steady-state flux distribution. The most interesting property of these EFMs is that their combinations (convex or linear depending on the existence of irreversibilities) can generate the complete flux space. Attending to this, the shared properties of the set of EFMs can be extrapolated to every possible flux distribution and thus, a proper analysis of these EFMs can lead to a better understanding of the capabilities of the related metabolic network.

## Results

### Genome-scale metabolic reconstruction

Metabolic reconstruction was based on the annotated pan-genome, derived from six previously published different strains of *Portiera* (ref. 14 and references therein). The pangenome contains a total of 280 genes, of which 160 were annotated as enzyme coding genes.A first draft of the metabolic network was generated using the SEED(16) pipeline. The draft obtained (id: Seed1206109.3.1165) included 136 internal reactions, 14 exchange fluxes and one biomass equation. Although the SEED pipeline includes a step for gap-filling the missing reactions, which are essential for the synthesis of all biomass precursors, the outputted model was not able to predict biomass formation. Moreover, ~95% of the reactions were found to be blocked. Thus, in order to obtain a consistent model, a detailed manual curation was performed using the unconnected modules approach(6) (see Supplementary Appendix 1 for details).

During the curation a total of 13 enzymatic activities, all related with amino acid biosynthesis and the pentose phosphate pathway (PPP) and coded by 10 genes, mostly contributed by the host(13–15), were added to the model (Table 1). To assess the essentiality of these orphan reactions, an *in-silico* knock-out analysis was performed. The result showed that only ribulose-phosphate 3-epimerase (EC 5.1.3.1) and glucose 6-phosphate isomerase (EC 5.3.1.9) do not significantly affect the predicted growth (see Supplementary Appendix 1, Figure 10, Table 1 labelled as *). In contrast, the synthesis of phosphoenolpyruvate (PEP) depends on either one of two orphan reactions: PEP synthase (EC 2.7.9.2) or, PEP carboxykinase (EC 4.1.1.49, Table 1 labelled as ^▴^), although the uptake of this metabolite from the host may not be discarded. Regarding the synthesis of branched chain amino acids, BCA (see Supplementary Appendix 1, Figures 6 and 7), the behaviour of the model does not change whether the endosymbiont or the host performs the transamination step. Finally, the last step of the histidine biosynthesis, catalysed by histidinol dehydrogenase (EC 1.1.1.23), may be carried by the host even though it is included in the endosymbiont network (see Supplementary Appendix 1, Figure 2). As a consequence of the manual curation, a flux consistent (*i.e*. every reaction can carry flux in the stationary state) genome-scale model of *Portiera*, named *i*JC86 predicting biomass formation was obtained (see Supplementary Table 1).

**Table 1:** Orphan reactions of the *i*JC86 model of *Portiera′s* metabolism. Reactions in bold could be substituted by a proper complementation with the host. Reactions labelled with a (*) are not essential for biomass production by *i*JC86. At least one of the reactions labelled with a (^▴^) must be active for biomass production by *i*JC86. BCA, branched-chain amino acid.

| REACTION | GENE | PATHWAY |
| --- | --- | --- |
| **histidinol dehydrogenase** | *hisD* | **histidine biosynthesis** |
| **histidinal dehydrogenase** | *hisD* | **histidine biosynthesis** |
| phosphoenolpyruvate synthetase ▲ | *pps* | chorismate biosynthesis |
| phosphoenolpyruvate carboxykinase ▲ | *pck* | chorismate biosynthesis |
| transaldolase | *talB* | pentose phosphate pathway |
| ribose-5P-isomerase | *rpiA/B* | pentose phosphate pathway |
| phosphogluconolactonase | *spontaneous/pgl* | pentose phosphate pathway |
| ribulose-5P-3-epimerase * | *rpe* | pentose phosphate pathway |
| glucose-6P-isomerase * | *pgi* | pentose phosphate pathway |
| **leucine transaminase** | *ilvE* | **BCA biosynthesis** |
| **isoleucine transaminase** | *ilvE* | **BCA biosynthesis** |
| **valine transaminase** | *ilvE* | **BCA biosynthesis** |
| **carbonic anhydrase** | *yadF/cynT/spontaneous* | **arginine biosynthesis** |

### Assessing the metabolic capabilities of *Portiera*

According to *i*JC86 *in-silico* predictions, *Portiera* is able to produce β-carotene and nine essential amino acids for the whitefly (see Table 2). Although the majority of enzyme-coding genes for these pathways have been found in the endosymbiont genome, some of the reactions are orphans. These orphan reactions include the transamination step at the end of the BCA biosynthesis, as well as the last two steps in the histidine pathway (Table 1). The stoichiometric analysis of *i*JC86 suggested that certain intermediate metabolites of these pathways must be exchanged between the endosymbiont and its host. In view of these observations, different scenarios of metabolic complementation are proposed (Figure 1). The first one is related to the exchange of glutamate and α-ketoglutarate. The whitefly provides glutamate to *Portiera*, which spends it on several transamination reactions. The *α*-ketoglutarate produced as a by-product can either be used as a precursor in the lysine biosynthesis pathway or transported to the insect. A second metabolic complementation event appears as a consequence of the by-product formation of 5-amino-1-ribofuranosyl-imidazole-4-carboxamide (AICAR) during histidine biosynthesis. Although AICAR is also a precursor in the *de novo* synthesis of purine bases, *Portiera* lacks the genes associated with this pathway. Accordingly, AICAR might be exported to the bacteriocyte cytoplasm, where it could be used by the host metabolism.

**Table 2:** Characteristics of the *i*JC86 model of *Portiera′s* metabolism. The first block displays the inputs and outputs needed for β-carotene and each amino acid to be produced. All the compounds in this block are essential for the whitefly. The second block contains those compounds (excluding ions), which are essential for the network to be functional and produce biomass. TPP, thiamine pyrophosphate; AICAR, 5-amino-1-ribofuranosyl-imidazole-4-carboxamide.

|  | INPUTS | OUTPUTS |
| --- | --- | --- |
| *leucine* | D-glucose, pyruvate/oxaloacetate, ADP | α-ketoglutarate, AMP |
| *valine* | D-glucose, pyruvate/oxaloacetate, ADP | α-ketoglutarate, AMP |
| *isoleucine* | D-glucose, pyruvate/oxaloacetate, aspartate, ADP | α-ketoglutarate, AMP |
| *tryptophan* | D-glucose, pyruvate/oxaloacetate, ADP, serine | α-ketoglutarate, AMP |
| *phenylalanine* | D-glucose, pyruvate/oxaloacetate, ADP | α-ketoglutarate, AMP |
| *threonine* | D-glucose, aspartate, ADP | α-ketoglutarate, AMP |
| *lysine* | D-glucose, pyruvate/oxaloacetate, aspartate, ADP | succinate, α-ketoglutarate, AMP |
| *arginine* | aspartate, glutamine, ornithine, ADP | fumarate, AMP |
| *histidine* | D-glucose, glutamine, ADP | AICAR, α-ketoglutarate, AMP |
| *B-carotene* | geranylgeranyl diphosphate | --- |
| **BIOMASS** | D-glucose, pyruvate/oxaloacetate | fumarate, succinate, α-ketoglutarate |
|  | alanine, proline, glycine, serine, glutamine, glutamate, asparagine, aspartate, cysteine, methionine, tyrosine, ornithine | --- |
|  | ubiquinone-8/menaquinone-8, lipoamide, coenzyme A, NAD<sup>+</sup>, NADP<sup>+</sup>, TPP | --- |
|  | nitrogenous bases and derivatives, lipids | AICAR, AMP |

**Figure 1:**
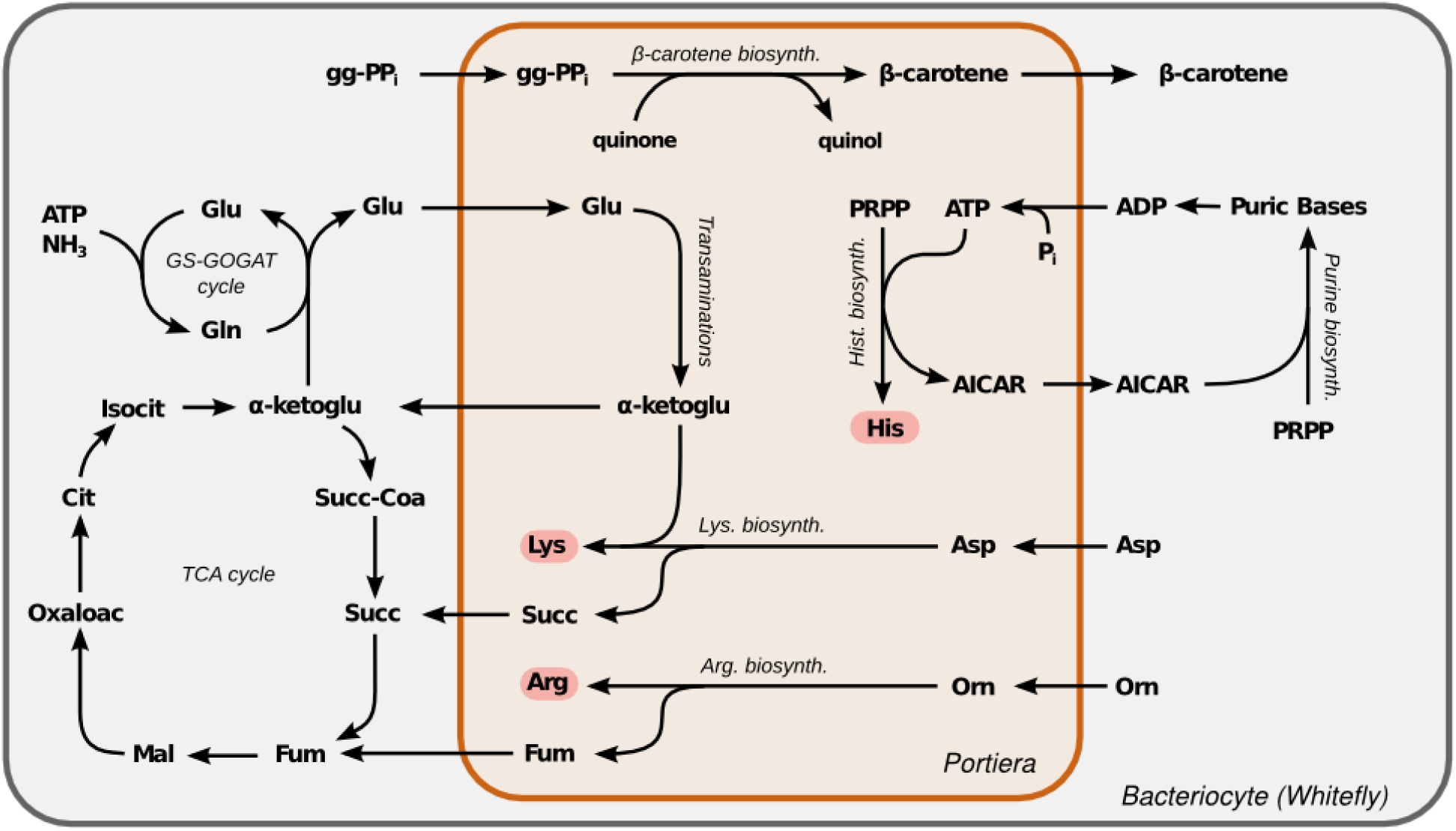
Metabolic complementations suggested by the *i*JC86 model. Exchange of glutamate and α-ketoglutarate: the endosymbiont uses glutamate for transamination reactions and the produced α-ketoglutarate is exported to the bacteriocyte. Exchange of AICAR and purine bases (and its derivatives): the endosymbiont generates AICAR as a subproduct of histidine biosynthesis and exports it to the bacteriocyte, where it is recycled into purine bases for the use of both the endosymbiont and its host. β-carotene production: whitefly provides geranylgeranyl diphosphate for the synthesis of β-carotene. This process requires the reduction of quinone to quinol, which is then oxidized by the respiratory chain. Exportation of succinate and fumarate: the synthesis of lysine and arginine involves the production of succinate and fumarate, respectively, compounds that must be consumed by the host. gg-PPi, geranylgeranyl diphosphate; Isocit, isocitrate; Cit, citrate; Oxaloac, oxaloacetate; Mal, malate; Fum, fumarate; Succ, succinate; Succ-Coa, succinyl-CoA; α-ketoglu, α-ketoglutarate; AICAR, 5-amino-1-ribofuranosyl-imidazole-4-carboxamide; PRPP, phosphoribosyl pyrophosphate; Orn, ornithine; GS, glutamine synthetase; GOGAT, glutamine: α-ketoglutarate aminotransferase.

The synthesis of β-carotene also involves metabolic collaboration between *Portiera* and the whitefly. According to the model, the biosynthetic precursor geranylgeranyl diphosphate should be imported from the bacteriocyte and reduced to β-carotene using ubiquinone which is reoxidized by a membrane-associated ubiquinol oxidase (EC 1.10.3.10), generating a proton-motive force.

The last complementation suggested by the model involves the bacteriocyte tricarboxylic acid (TCA) cycle, a pathway absent in the endosymbiont. In *Portiera*, the synthesis of lysine and arginine entails the production of succinate and fumarate, respectively. However, since the endosymbiont lacks any reaction able to consume these compounds, a plausible scenario is that both metabolites are exported to the bacteriocyte’s cytoplasm.

### Dissecting *Portiera’s* energetics through elementary flux modes analysis

One remarkable feature of *Portiera′s* metabolism is that it lacks several canonical energy-conserving pathways, namely, glycolysis, a complete TCA cycle, and fatty-acid oxidation. Furthermore, only a fragmented PPP, and the production of succinyl-CoA and acetyl-CoA are retained in the endosymbiont. This striking characteristic is shared with other endosymbionts of equal or smaller genome size (*e.g. Carsonella, Sulcia, Uzinura, Tremblaya* and *Hodgkinia*, see(10)). Because *Portiera* can synthesize the same number of amino acids as endosymbionts whith larger genomes, the characterization of its energetics could serve as a model for the study of the endosymbionts harbouring the most reduced genomes whith equal or more limited biosynthetic capabilities.

A detailed study of the energetics of *Portiera* was performed by analysing the elementary flux modes (EFMs) of the *i*JC86 model (see Methods). Briefly, an EM contains a minimal and unique set of enzymatic reactions that can support cellular functions at steady state and, more important, any metabolic behaviour under this assumption can be expressed as combination of these EMs. Thanks to this property, the evaluation of the EFMs associated with the production of energy, referred to as energy-producing EFMs (enEFMs, see Elementary Flux Modes Analysis in the Methods section), allows the characterization of every energetic pathway in the model.

***Definition I:** an energetic EFM (**enEFM**) of a metabolic network is an EFM that produces a net amount of energy in the form of ATP.*

The *i*JC86 model accounts for 6,503 EFMs, of which only 34 are enEFMs. To completely characterise the energetics associated to these enEFMs, each one was assigned with seven properties: net NADPH production; net NADH production; H+gap (net cytosolic H^+^ consumption not considering the cytosolic H^+^ lost by pumping reactions); ATP consumed; net ATP production; fraction of produced ATP corresponding to the H^+^ gap; and the number of transported molecules through the membrane (see Supplementary Table 2 for a complete list of the enEFMs and their properties). The relation between these properties is deduced and discussed in Supplementary Appendix 2.

Essentially, every production in the network is either an amino acid or it is linked to the synthesis of one. Based on this, it is possible to characterize each enEFM through the assignment of an order, which is defined as follows:

***Definition II:** the **order** of an enEFM is the number of amino acids that are produced in this enEFM.*

Furthermore, the amino acids produced by *Portiera* can be classified as follows:

***Definition III:** an amino acid is energetic if it is produced in an order one enEFM. In general, a set of amino acids is called energetic if, and only if, no other enEFM produces a subset of these same amino acids.*

***Definition IV:** an amino acid is defined as energy-dependent if, after discounting the contribution of the energetic amino acid sets, the amount of energy produced in each particular enEFM involved is less than zero (energetically self-sufficient if equal to zero).*

It is important to consider that, in general, these definitions hold only if no zero order enEFMs exist in the model: if the former is not true, it cannot be assured that the energy produced in enEFM of order > 0 is related to the amino acids synthesized in it. However, the *i*JC86 model has one. This enEFM comprises solely the synthesis of β-carotene from the geranylgeranyl diphosphate imported from the host (see Supplementary Appendix 1, Figure 15). In this pathway four double bounds are formed, each of which requires the concomitant reduction of a quinone to quinol. These four quinol molecules are incorporated and reoxidized in the respiratory chain, and 2.5 molecules of ATP *per* β-carotene synthesized may be produced. Because no other enEFM is associated with the production of β-carotene, it can be assumed that definitions III and IV hold even with the presence of this zero order enEFM. The characteristics of the enEFMs, which only produce energetic amino acid sets, are given in Table 3, and a representation of the energetics of *Portiera* is displayed in Figure 2.

**Figure 2:**
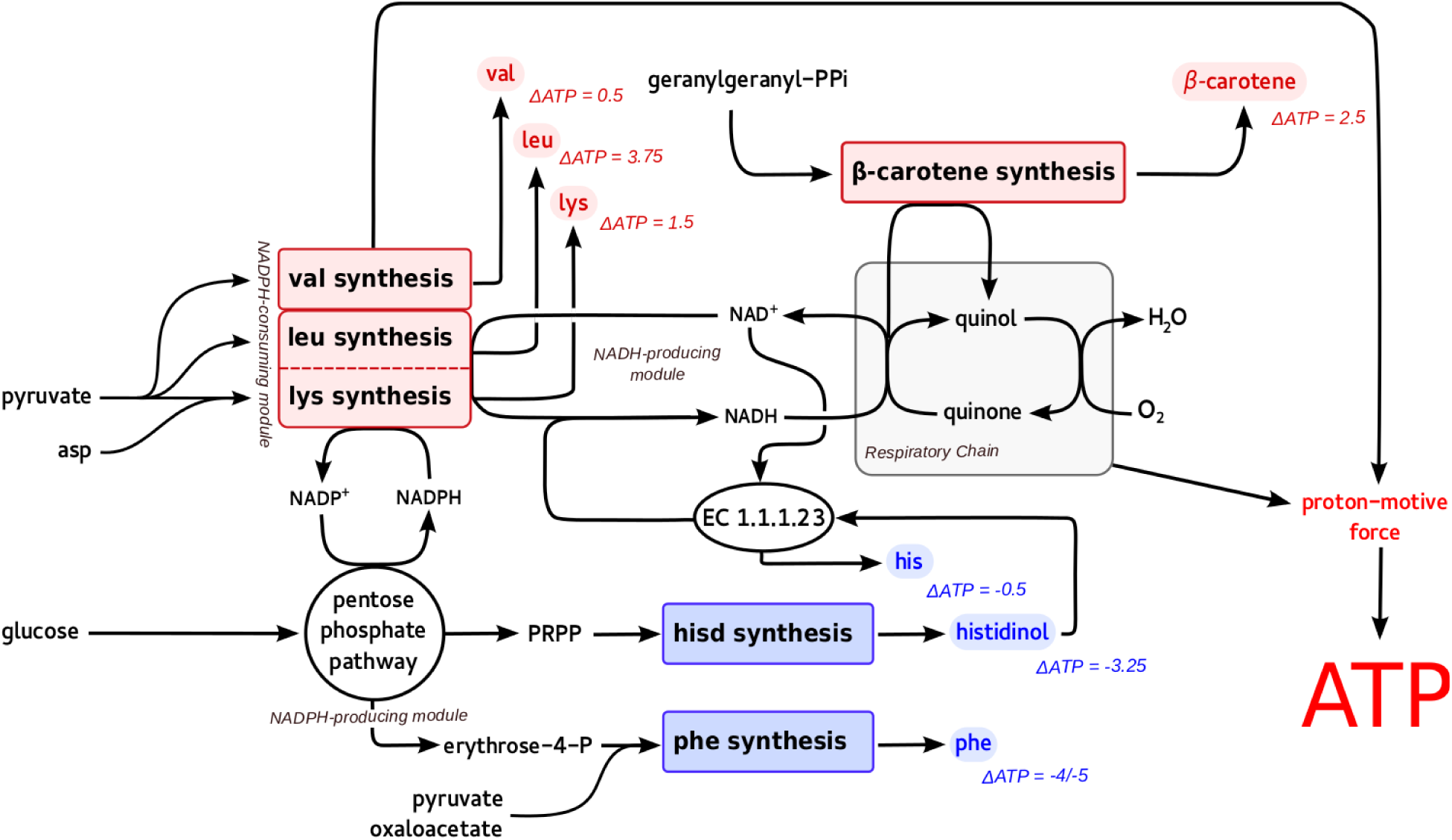
Representation of the energetics of *Portiera* according to the *i*JC86 model. The only zero order enEFM of the *i*JC86 model involves the synthesis of β-carotene. In this pathway the geranylgeranyl diphosphate is oxidized by quinones, which are then incorporated into the respiratory chain in the form of quinol, generating ATP through oxidative phosphorylation. Every non-zero order enEFM is associated with the production of at least two amino acids, one associated with the production of NADPH (phenylalanine or histidine/histidinol) and the other with its consumption (leucine, valine or lysine). The number of ATP molecules generated (red) /consumed (blue) *per* molecule of amino acid or amount of β-carotene produced is represented by the thickness of the arrows. Dotted arrows represent pathways in which ATP synthesis is associated only with the gradient generated through the consumption of intracellular protons. PRPP, phosphoribosyl pyrophosphate.

**Table 3:**
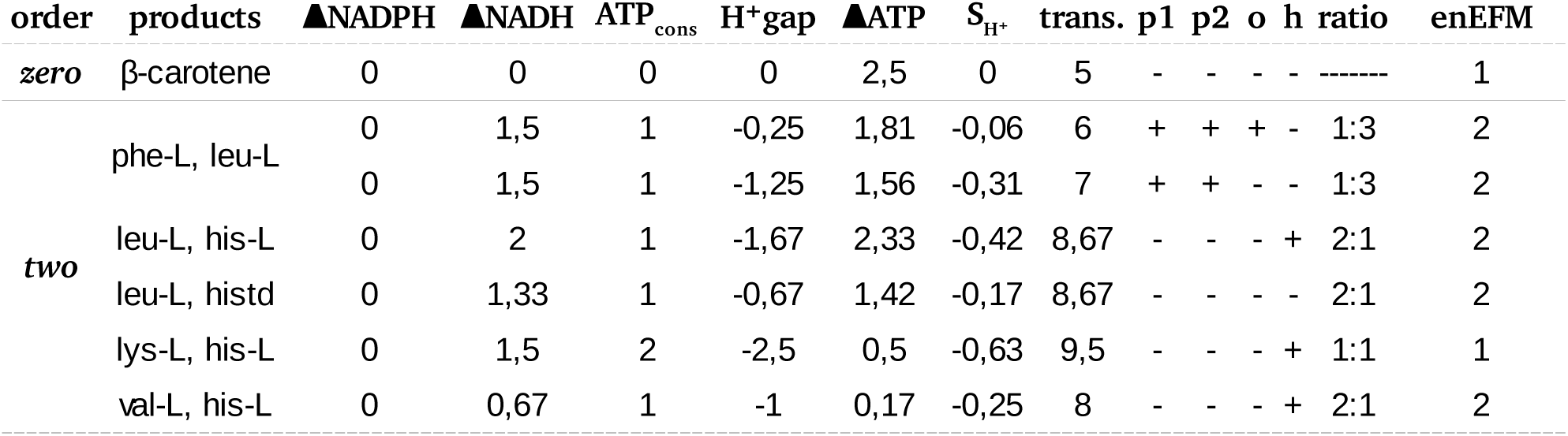
Properties of the EFMs related to the energetic productions of the *i*JC86 model *Portiera*. For each order (first column), the corresponding amino acid sets (β-carotene for orde zero) are displayed in the second column, while the associated properties are listed in the next ones. The properties listed are: net NADPH production ΔNADPH); net NADH production ΔNADH); ATP consumed (ATP_cons_); proton gap (H_+_gap); net ATP produced ΔATP); ATP produced due to the proton gap (S_H_^+^); and number of transported molecules (trans). Data are based on the production of 1 arbitrary flux unit of amino acid. The requests for each enEFM regarding the presence (+) or absence (–) of a variant-specific reaction are indicated as follows requirement of D-ribulose-5-phosphate epimerase (p1) or D-glucose-6-phosphate isomerase (p both defining the *i*JC86 *p-* model; oxoaloacetate-dependent phosphoenolpyruvate production ( requirement of histidinol oxidoreductase (h). The number of enEFMs associated with each energetic amino acid set and each particular property are displayed in the last column. For eve second order EFM the contribution of each amino acid is displayed (see Ratio column). The contribution of β-carotene biosynthesis was discounted in every EFM, which relies on it. Histc histidinol.

| order | products | ΔNADPH | ΔNADH | $ATP_{cons}$ | $H^+$gap | ΔATP | $S_{H^+}$ | trans. | p1 | p2 | o | h | ratio | enEFM |
| --- | --- | --- | --- | --- | --- | --- | --- | --- | --- | --- | --- | --- | --- | --- |
| *zero* | β-carotene | 0 | 0 | 0 | 0 | 2,5 | 0 | 5 | - | - | - | - | ----- | 1 |
| *two* | phe-L, leu-L | 0 | 1,5 | 1 | -0,25 | 1,81 | -0,06 | 6 | + | + | + | - | 1:3 | 2 |
|  |  | 0 | 1,5 | 1 | -1,25 | 1,56 | -0,31 | 7 | + | + | - | - | 1:3 | 2 |
|  | leu-L, his-L | 0 | 2 | 1 | -1,67 | 2,33 | -0,42 | 8,67 | - | - | - | + | 2:1 | 2 |
|  | leu-L, histd | 0 | 1,33 | 1 | -0,67 | 1,42 | -0,17 | 8,67 | - | - | - | - | 2:1 | 2 |
|  | lys-L, his-L | 0 | 1,5 | 2 | -2,5 | 0,5 | -0,63 | 9,5 | - | - | - | + | 1:1 | 1 |
|  | val-L, his-L | 0 | 0,67 | 1 | -1 | 0,17 | -0,25 | 8 | - | - | - | + | 2:1 | 2 |

While no enEFMs of order one has been found, a total of eleven enEFMs of order two were identified. These enEFMs involve four distinct pairs of amino acids (see Table 3): leucine/histidine; valine/histidine;lysine/ histidine; and phenylalanine/leucine. Of the three sets containing histidine, only the leucine/histidine pair generates energy in the absence of the histidinol oxidoreductase activity. In addition, only the amount of ATP generated by the synthesis of leucine/histidine is significant: 2.33 mols of ATP (1.42 if the histidinol oxidoreductase is discarded) *per* mol produced of histidine/leucine versus 0.17 in the case of valine/histidine and 0.5 in the case of lysine/histidine. This suggests that only the set involving histidine and leucine can be considered energetic with reasonable certainty. The possibility of a certain amount of active transport, together with the high number of transport events required, suggests that the other histidine-containing pairs are probably not energetic but, at least, cheap to produce.

Since the synthesis of phosphoenolpyruvate (essential precursor in aromatic amino acid production pathways) relies on either pyruvate or oxaloacetate, the four phenylalanine/leucine enEFMs can be classified into two groups: pyruvate-dependent or oxaloacetate-dependent. The reactions involved in each case are:

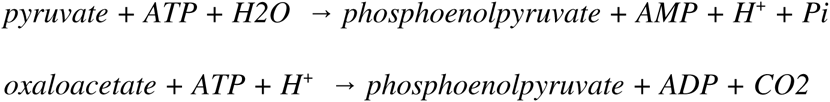

Because the *i*JC86 model assumes that the recycling of AMP to ADP depends on the host, regarding energetics there are two significant differences between the two groups of phenylalanine/leucine enEFMs. Firstly, the usage of pyruvate involves more transport processes compared to oxaloacetate due to the recycling of AMP in the insect cytoplasm. Secondly, while the phosphoenolpyruvate synthesis via pyruvate generates one cytosolic H^+^, if oxaloacetate is used instead, one cytosolic H^+^ is consumed. These differences explain why distinct enEFMs were identified, since the usage of oxaloacetate/pyruvate implies different reactions.

Another important property of the phenylalanine/leucine producing enEFMs is that they all depend on the presence of two non-essential orphan reactions. The absence of either D-glucose-6-phosphate isomerase or D-ribulose-5-phosphate-3-epimerase supposes the coupling of phenylalanine and tryptophan production (Figure 3). Because tryptophan is an expensive amino acid to produce in the network, this coupling could seriously restrict not only the phenylalanine supply to the whitefly, but also the energetic effects of the endosymbiont. If we focus on the viability of the enEFMs studied above, oxalacetate-dependent ones produce 1.81 ATP *per* mol of amino acid while pyruvate-dependent ones produce 1.56 ATP *per* mol of amino acid. In this case the difference is slight, and both enEFMs can be considered energy-producing pathways.

**Figure 3:**
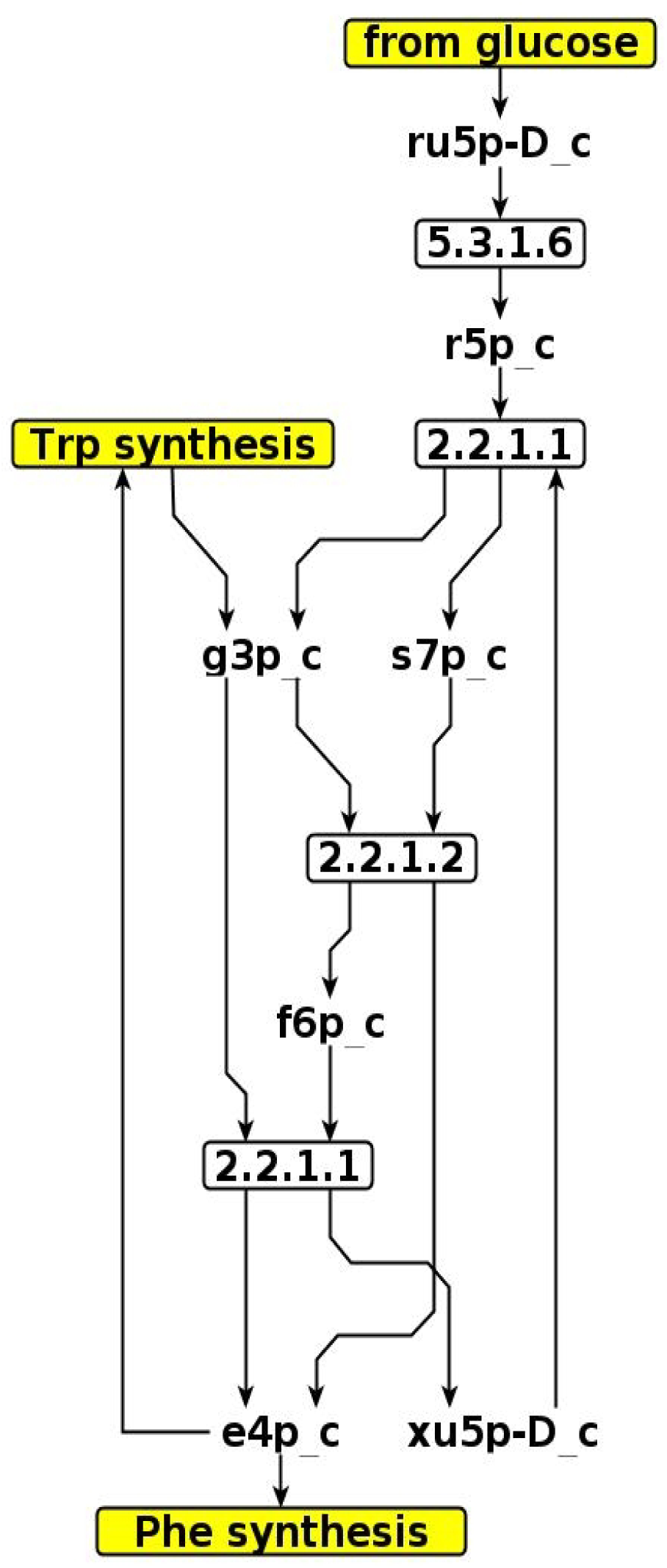
Graph representing the coupling between phenylalanine and tryptophan biosynthesis in *i*JC86 model of *P. aleyrodidarum* metabolism if D-ribulose-5-phosphate-3-epimerase and D-glucose-6-phosphate isomerase are not included as orphan. Metabolites are represented with their abridged name and reactions are represented as softened rectangles containing the corresponding EC number. ru5p-D_c, ribulose-5-phosphate; g3p_c, glyceraldehyde 3-phosphate; s7p_c, sedoheptulose-7-phosphate; f6p_c, fructose-6-phosphate; e4p_c, erythrose-4-phosphate; xu5p-D_c, xylulose-5-phosphate.

For the characterization of the order two enEFMs to be complete, the contribution of the synthesis of each amino acid to the energy production must be identified. As can be seen in Table 3, the net production of NADPH is zero in each of these enEFMs. Because leucine, valine and lysine consume NADPH in their synthesis, the other amino acid must produce the same amount of reduced cofactor. In order to take advantage of this situation, a reaction that freely interchanges NADPH and NADP^+^ was included in the model to obtain the energetic contribution of each amino acid in every energetic pair: if the synthesis of a particular amino acid in the pair produces ATP, new order one enEFMs must appear. The reason is that the inclusion of the NADP^+^-NADPH interchanging reaction buffers the NADPH consumed or generated during the synthesis of these amino acids and thus, it does not need to be obtained /redirected from another route. This analysis shows that leucine, valine and lysine are the ATP generators in the energetic pairs while histidine (or histidinol) and phenylalanine provide NADPH while consuming part of this ATP (Supplementary Table 3). As expected, leucine and lysine obtain ATP through the integration of NADH into the respiratory chain. However, no NADH is produced in the valine synthesis, with the ATP produced coming solely from the H^+^ gap. This means that the production of valine consumes cytosolic H^+^ that is excreted as part of other molecules (*e. g*. water), thereby increasing the electrochemical potential of H^+^ and allowing ATP synthase to operate. This effect is present in other enEFMs, although this is the only mode in which this effect is the unique source for ATP synthesis.

With respect to higher order enEFMs, the amino acids produced by each of these EFMs always include one of the amino acid pairs synthesized by second order enEFMs. This means that, by definition, there is no energetic amino acid sets with more than two components in any of the *i*JC86 variants considered.

Although it is true that only pairs of amino acids were classified as energetic, it is possible that, when produced together or with other amino acids in the same enEFM, the flux distribution allows a higher production of energy than that expected from the order two enEFMs. These differences may be due to the utilization of a different pathway in the production of the energetic amino acids. However, when discounting the contribution of β-carotene and energetic amino acids synthesis, no such effects were observed. Finally, the case of energetically self-sufficient EFMs which produce amino acids was considered. However, no EFM matching this criterion was found.

### The role of β-carotene synthesis in the metabolic capabilities of *Portiera*

One remarkable feature of *Portiera*’s metabolism involves the ATP production associated with β-carotene synthesis. It is worth to note that the turnover of this molecule may not be high enough to support *Portiera’s* energetic needs. If this was the case, the remaining ATP requirements should be satisfied by a non-negative combination of non-zero order enEFMs. This implies that, in order to growth, the endosymbiont may overproduce certain amino acids. To study this phenomenon the maximum biomass produced and the magnitude and nature of the overproductions were determined after constraining the flux through β-carotene synthesis to different values (from zero to unbounded). Because the majority of the amino acid production may be destined to the insect, a reaction representing the whitefly biomass was included in the model and several ratios of endosymbiont biomass to insect biomass were evaluated (see Methods). Since no specific information on the amino acid composition and ATP requirements of either the whitefly or *Portiera* was found in the literature, the simulations were performed using an ensemble of randomized biomass equations derived from those corresponding to *A. pisum* and *E. coli*, respectively (see Methods). It is important to note that the ATP requirement for the whitefly biomass equation in *i*JC86 is expected to be zero.The reason for this expectation is that the cost of assembling the proteins needed for the insect is assumed by the insect itself and thus, is not included in the model. Thus, the ATP cost of the biomass equations of the whitefly was set to be zero (see Methods). The most relevant findings obtained through this analysis are displayed in Figure 4.

**Figure 4:**
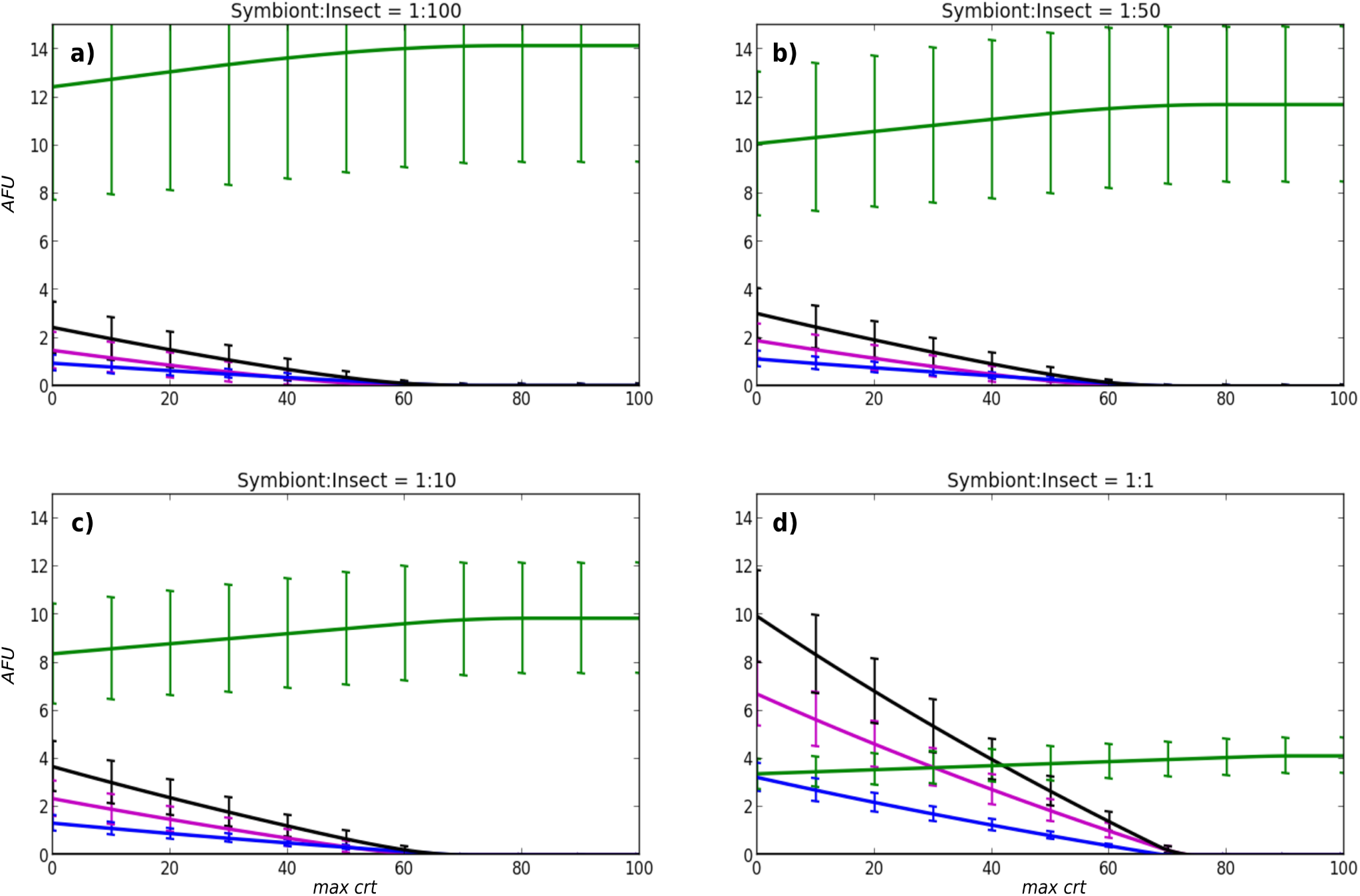
Impact of β-carotene synthesis on the metabolic performance of the *i*JC86 model of *Portiera’s* metabolism. Representation of the effect of β-carotene synthesis in maximum insect biomass production (green) and amino acid overflows (histidine, blue; leucine, magenta; histidine + leucine, black). Because part of the insect’s biomass corresponds to *Portiera*, several symbiont-to-insect biomass ratios were tested (panels *a, b, c* and *d*). For each ratio, the 90% of maximum biomass production, in addition to the overflow of each produced amino acid were calculated under different constraints on the β-carotene synthesis (*max crt*): from 0% to 100% of the total carbon sources flux. Only leucine and histidine were detected to be overproduced. Due to the lack of specific information regarding the biomass equation of either *Portiera* or the whitefly, the simulation described above was carried on an ensemble of 100 randomized variants of these equations (see *Methods*). The results plotted here accounts for the mean of all simulations, error bars representing the corresponding standard deviation. *AFU:* arbitrary flux units.

Because the differences in the amino acid compositions of the two biomasses are less marked than the corresponding to ATP consumption, it can be assumed that an increase in the endosymbiont to insect biomass ratio is equivalent to an increase in the ATP requirements for the whole system (whitefly plus *Portiera*) to grow. Thus, attending to the results presented in the previous section, an increment in the overproduction of amino acids is expected. Results from constraining the flux through β-carotene synthesis demonstrate this point: as the ratio of endosymbiont biomass to insect biomass is increased, the total biomass produced (insect plus endosymbiont) quickly decreases at the same rate as the flux of overproduced amino acids increases (Figure 4).

The effect on biomass production seems to be independent of the constraint in β-carotene synthesis but not for the amino acid overflows (Figure 4). If the β-carotene production accounts for at least around 60% of the total carbon source flux, no amino acid overflow is observed irrespective of the value of the symbiont-to-insect biomass ratio. This suggests that, assuming the most relaxed case (symbiont-to-insect biomass ratio = 1:100 in the model with all the orphan reactions of Table 1 included), the number of β-carotene molecules produced *per* amino acid destined to biomass is at least 2. Because a minimum of 90% of the biomass maximum was set in these analyses, we asked at what fraction of the biomass maximum (if this maximum exists) will there be no β-carotene production and no amino acid overflow. Restricting the β-carotene and amino acid exchange fluxes to zero impedes the growth of the system, whatever the symbiont-to-insect biomass ratio. Even if this condition is relaxed (by allowing a maximum flux through these reactions of 1% of the total carbon source flux allowed to enter the symbiont), less than 10% of the maximum biomass is produced in the corresponding models without these constraints. Taking these results as a whole, for *Portiera* (no matter which orphan reactions are included in the model, see Table 1) to work in near-optimal (>90% of the maximum, as above assesed) biomass-producing conditions (either for itself or for the insect) it must either overproduce amino acids or β-carotene.

If we consider the nature of the overproduced amino acids (Figure 4), it is not surprising that leucine and histidine/histidinol are the ones which are overproduced. As mentioned above, the leucine-histidine/histidinol and phenylalanine-leucine pairs are associated with the most effective second order enEFMs, with respect to the amount of ATP synthesized. Because leucine participates in all the pairs named, it is overproduced in higher amounts than histidine/histidinol. Phenylalanine is not overproduced essentially because it is required in higher amounts in the biomass equations compared to histidine, and because the leucine-histidine pair produces approximately 0.5 ATP molecules *per* amino acid, which is more than the best phenylalanine-leucine-producing enEFM.

## Discussion

The results of the stoichiometric analysis of the *i*JC86 model with regard to the amino acid metabolism are consistent with all previous studies on metabolic complementation between *Portiera* and its insect host. In addition, these complementations resemble those reported in *Buchnera aphidicola*, the endosymbiont of the aphid *Acyrthosiphon pisum(17, 18)*. Most of these studies are based either on qualitative genome analysis(11, 12, 14) or on experimental approaches (e.g. transcriptome analysis(13, 15)). A curated model, such as the here presented *i*JC86, offers additional opportunities for investigating the structural properties of the metabolic network whith a systems biology approach, such as the EFM analysis. According to the *i*JC86 model, the energetics of *Portiera* relies (whith the exception of the spontaneous proton-motive force generated in the synthesis of valine) entirely on its respiratory chain: all the ATP produced comes from either the oxidation of the ubiquinol produced in the formation of the double-bonds of the β-carotene or the oxidation of the NADH generated during the synthesis of certain amino acid pairs (Figure 2). These pairs are established because the NADPH balance must be maintained. In particular, they involve a NADPH-consuming/energy-producing amino acid (leucine, valine or lysine) and a NADPH-producing/energy-consuming one (phenylalanine, histidine/histidinol). The remaining amino acids (arginine, threonine, tryptophan and isoleucine) are all energy dependant. The amount of energy produced in association with the synthesis of these amino acids ranges from 0.17 to 2.33 moles of ATP *per* mol of amino acid produced (Table 3). However, these results only hold if all the transport processes in *Portiera* are assumed to be passive. Although this assumption is reasonable considering the high concentration of sugars and non-essential amino acids in the insect diet(19), it may not be wholly valid. Minor deviations from the allpassive transport assumption has important consequences when considering the energetics of *Portiera*. Essentially, the ATP production associated with the valine/histidine and lysine/histidine pairs would be abolished, leaving β-carotene and leucine production as the energetic motor of the symbiont. This situation highlights the importance of the knowledge gap regarding the transport processes in endosymbionts when analyzing their metabolic networks: the behavior of the network can vary dramatically under slightly different scenarios. Nevertheless, the passive transport assumption allows all possible functionalities of the network to be considered, and robust behaviors can be identified, such as the case of β-carotene or the leucine pairs discussed previously.

Two remarkable features were revealed through the modelling of *Portiera*’s energetic metabolism. The first one comprises the production of ATP associated with valine synthesis (Supplementary Table 3). Because no substrate-level phosphorylation exists in the network, an energy production independent of the respiratory chain was unexpected, and raises the question of how the endosymbiont can generate the electrochemical proton gradient needed for the ATP synthase to operate. According to the *i*JC86 model predictions, the proton motive force needed is obtained through the exportation of cytosolic protons by including them in the structure of other molecules. Symmetrically, this contribution to the generation of proton motive force, defined here as a proton gap, can also be detrimental to energy production if protons are generated instead of consumed. The effects of the proton gap on the symbiont energetics are widespread and significant in the majority of cases (SupplementaryTable 3).Because this effect is only predicted using stoichiometric analysis, empirical tests should also be used to assess its role in the system.

A second remarkable feature of *Portiera*’s metabolism involves the β-carotene synthesis. Carotenoids and their derivatives have important functions in insects, such as coloration, vision, diapause, photoperiodism, mate choice or oxidative stress(20). The stoichiometric analysis of the *i*JC86 model reveals an additional possible function related to energetics and, ultimately, to the control of amino acid biosynthesis. Leucine, lysine, valine and, to a lesser extent, phenylalanine and tyrosine (obtained through phenylalanine oxidation), span a notable percentage of the proteome not only in insects(21) but also in other organisms(22). In contrast, histidine is only required in small quantities(21, 22). If no β-carotene-associated ATP production exists, *Portiera* will produce an excess of certain amino acids with a significant overproduction of leucine and histidine (Figure 4). However, if β-carotene-associated ATP production is allowed, this restriction can be avoided. Nevertheless, the minimum amount of β-carotene synthesis needed for neglecting amino acid production is about two times the production of amino acids, if all of them are destined to the whitefly, and much more if we also consider *Portiera’s* growth. Thus, for the insect-endosymbiont system to growth whith a significant yield (>10% of its maximum), either leucine and histidine or β-carotene must be substantially overproduced. This performance suggests two non-exclusive possible scenarios: the overproduced substances have a particular physiological or ecological (*e.g*. through excretion in honeydew) function for the whitefly, or they have no significant fitness effect in the system. However, a third scenario can be conceived taking into consideration recent evidence(23) which suggests that β-carotene could be involved in light-induced reduction of NAD^+^ in insects. Thus, if light can induce a ciclic electron transport in which the NADH oxidation through NADH-dehydrogenase takes part, ATP can be synthesized whithout the concomitant production of amino acids or β-carotene. Assuming this as true, its energetic role in *Portiera* is independent of the amount of β-carotene synthesized, and thus overproduction of any kind is unexpected.

It is worth to emphasise that the properties inferred from *i*JC86 model of *Portiera’s* panmetabolism can be extrapolated to similar networks with few additional considerations. Lets consider two examples. Attending to comparative genomics(14), the only *Portiera* strain which deviate significantly from the others is *Candidatus* Portiera aleyrodidarum BT-QVLC, hosted in *B. tabaci*. Specifically, this strain lacks genes involved in lysine (*dapB*, *dapF* and *lysA*) and arginine (*argG* and *argH*) synthesis(14, 15). The inspection of the reactions associated to *dapB, dapF* and *lysA* shows that they contribute with 0.25 ATP *per* amino acid (by increasing the proton gap by one *per* amino acid) to the net production of 0.5 ATP *per* amino acid associated to the enEFM’s in which lysine is produced (Table 2). Because arginine synthesis is not associated by any means to ATP or NADPH production, it can be concluded that the energy metabolism of *Candidatus* Portiera aleyrodidarum BT-QVLC is essentially the same than the one inferred from *i*JC86 model of *Portiera’s* panmetabolism. Let consider now the effects on the network of allowing the importation of phosphoenolpyruvate (PEP) in *Portiera’s* energy metabolism. The only destination of this metabolite in the network is the production of phenylalanine. Attending to Supplementary Table 3, the cost of synthesizing this amino acid depends, precisely, on the precursor employed in the production of PEP.Thus, by discounting the contribution of PEP synthesis to those enEFM associated to phenylalanine production, the effects of the free importation of PEP on the energy metabolism of *Portiera* can be determined. In particular, a new order two enEFM synthesizing phenylalanine/leucine will appear with an associated net production of ~2.06 ATP *per* amino acid, in contrast to the ~1.86 and ~1.56 ATP obtained if PEP is produced from oxaloacetate and pyruvate, respectively. Again, no substantial changes appear if a free incorporation of PEP is considered, the main reason being that the more efficient ATP production in the network is still associated to the synthesis of β-carotene (2.5 ATP *per* β-carotene) and the pair leucine/histidine (~2.33 ATP *per* amino acid).

In conclusion, *Portiera* shows the capability of simultaneously providing the whitefly with nine essential amino acids while being capable of synthesizing its own ATP. The ways nature has allowed this to be possible involve the efficient configuration of biosynthesis pathways that reduces the catabolic reactions needed to a minimum. This configuration is especially notable when β-carotene synthesis is considered: just a few reactions allow a coarse control of amino acid production while serving as fuel for this particular ATP-synthesis machinery. Because *Portiera* is just one of a number of reduced genome endosymbionts, it serves as a milestone in the exploration of the energetics of these organisms.

## Methods

### Elementary Flux Modes Analysis

Computation of the elementary modes (EFMs) was performed on a modified version of *i*JC86, *where* an ATP consumption reaction (referred to as *energy*) was incorporated as a label for the identification of enEFMs. EFMs in which *energy* is greater than zero are the enEFMs of the model (see *definition I* in results). The set of EFMs was calculated using the EFMtool(24).

### Determination of the impact of β-carotene synthesis on the metabolic performance of *i*JC86

In order to perform this analysis the model was reconfigured to represent the *Portiera-* whitefly interaction in a more realistic way. First, a simplified biomass equation representing the amino acid demands required for the insect’s growth was added to the system, using the amino acid content of *A. pisum* as a reference(21). In particular, each amino acid produced by *Portiera* was added as a reactant with a stoichiometric coefficient equal to its content in *A. pisum* (with the exception of phenylalanine, whose coefficient also includes the content of tyrosine because the last one is a necessary precursor of the first one). Once the stoichiometric coefficients were set, they were normalized for their summation to be one.

Due to the fact that the relation of both biomass productions is unknown, several symbiont-to-insect biomass ratios were tested. For each of these ratios, the maximum biomass production of the system (insect+symbiont) was calculated through flux balance analysis(25), after constraining the β-carotene exportation flux to several values *C* ∈ [0, 100]. Then, for each *C*, the overproduction of every amino acid was evaluated by considering the minimum value computed using flux variability analysis(24) (biomass flux >90% of its maximum).

To fulfil the previous calculations, the model’s reactions were considered as unbounded by setting their upper bounds to 10^6^ arbitrary flux units (AFUs) and their lower bounds to 0 or -10^6^ AFUs for reversible reactions. Moreover, a constraint to limit the maximum carbon sources (CS) importation was incorporated:

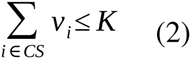

where *CS* includes glucose, pyruvate, oxaloacetate, α-ketoglutarate, ornithine, aspartate, and geranylgeranyl diphosphate; *v_i_* is the flux through the corresponding transportation reaction and *K* is the value of the constraint. The value of *K* was fixed to 100 AFUs in order to make it the only active constraint in the model.

Since no specific information on the amino acid composition of either the whitefly or *Portiera* was found in the literature, the experiments were performed using an ensemble of 100 randomized variants of the biomass equations to make the results obtained more robust. The randomization was conducted in the following way: for each amino acid the stoichiometric coefficient was drawn from a uniform distribution in the interval (0.75 *x*_0_, 1.25*x*_0_) where *x*_0_ corresponds to the reference value. These values were normalized afterwards as mentioned above. The ATP consumption for the insect and symbiont production was then modified. For the insect, the ATP cost was set to zero. On the other hand, the ATP cost associated to the symbiont biomass equation was extracted from a uniform distribution in the interval [30,70].

### Computational Tools

Constraint-based analysis was performed using the python-based toolbox COBRApy(26). LP and MILP problems were solved using the Gurobi Solver(27) accessed through COBRApy. The computation of connected components for the detection of the set of unconnected modules was performed using the Python NetworkX library(28). The scripts developed for this paper were programmed in Python language(29), and can be requested from the authors. Graphs were drawn using the yEd Graph Editor(30). All the computations were performed on a workstation using an Intel® Xeon® CPU X5660 2.80GHz processor,with 32GiB, running under Ubuntu 14.04.2 Linux OS.

## Acknowledgements

We would like to thank the Obra Social Programme of *La Caixa* Savings Bank for the doctoral fellowship granted to JCE. Financial support from the Spanish Government (grant reference: BFU2012-39816-C02-01 and BFU2012-39816-C02-02, co-financed by FEDER funds and the Ministry of the Economy and Competitiveness) and the Regional Government of Valencia (grant reference: PROMETEOII/2014/065) is also gratefully acknowledged.

## Contributions

F.J.S. and D.S.G. provided functionally annotated genomic data prior to publication; J.C.E. and M.P.L. performed the experiments; J.C.E, M.P.L, F.M and J.P. analyzed the data; J.C.E. wrote the manuscript with inputs from M.P.L, F.M., D.S.G., F.J.S. and J.P. All authors read and approved the final manuscript.

